# Characterization of the role of several COPI complex isoforms during the early acceptance of compatible pollen grains in *Arabidopsis thaliana*

**DOI:** 10.1101/427534

**Authors:** Daniel A. Cabada Gomez, M. Isabella Chavez, Emily Indriolo

**Author notes:** Corresponding author: Emily Indriolo.

## Abstract

COPI is a seven subunit coatomer complex, consisting of α, β, β′, γ, δ, ε, and ξ; in *A. thaliana*, COPI is necessary for retrograde transport from the Golgi to the Endoplasmic Reticulum, Golgi maintenance, and cell-plate formation in plant cells. Vesicle recruitment to the pollen contact point is required for pollen hydration and pollen tube penetration. To determine what other aspects of trafficking may be involved in the stigmatic papillae acceptance of compatible pollen, knock-out lines of several isoforms of the COPI complex were characterized in their roles during compatible pollination. Isoforms that were studied included α1-COPI, β-COPI, β′-COPI, γ-COPI and ε-COPI. Each mutant line was characterized in regards to pollen grain adherence, pollen tube penetration, and seed set. Of the mutant lines examined, *α1-copi* had the strongest phenotype with issues with compatible pollen grain adherence, tube germination and reduction in seed set while other lines had milder but visible retardation in compatible pollen acceptance. The data presented here are the first study of the role of the COPI complex in compatible pollinations and that certain subunit isoforms are required for compatible pollen acceptance.

## Introduction

A critical point during the flowering plant life cycle, is the dynamic interplay between the pollen grain and the pistil, as the pistil regulates the acceptance or rejection of a pollen grain. In the mustard family of plants, the Brassiaceae, the stigma surface is a dry type, where the pollen grain must acquire water for germination and pollen tube growth from the unicellular stigmatic papillae cells (Heslop-Harrison 1979). Upon perception of the pollen grain by the stigmatic papilla, rapid communication occurs between these two cells, and the basal compatible pollen response is activated. At this instance, the stigma is able to reject foreign pollen and to begin the process of accepting the compatible pollen grain (for review, see (Chapman and Goring 2010; Dickinson 1995; Doucet et al. 2016; Hiscock and Allen 2008; Indriolo et al. 2014). The initial interaction between the exine of the pollen grain and the stigma surface is well characterized. If the pollen grain is compatible, it is able to form a lipid-protein interface known as a ‘foot’ and it is able to increase adherence to the papilla cell (Elleman CJ 1990; Elleman CJ 1992; Gaude and Dumas 1986; Mattson 1974; Mayfield et al. 2001; Preuss et al. 1993; Stead AD 1980; Zinkl et al. 1999). During foot formation, it is hypothesized that the Brassicaceae-specific compatible pollen recognition occurs, although the full mechanism is not yet clearly understood (Hulskamp et al. 1995; Pruitt 1999). There are two stigma-specific *Brassica oleracea* glycoproteins that play a role in this; S-Locus Glyco-protein and S-Locus Related1 (SLR1) (Luu et al. 1997; Luu et al. 1999) which may interact with two pollen coat specific proteins, PCP-A1 and SLR1-BP, respectively (Doughty et al. 1998; Takayama et al. 2000; Wang et al. 2017). At this same moment, self-incompatible pollen grains can be recognized and deactivation of the basal pollen response beings while simultaneously activating the self-incompatibility signaling pathway (for review see, (Dresselhaus and Franklin-Tong 2013; Indriolo et al. 2014; Sawada et al. 2014).

One of the most important actions that comes from the compatible pollen recognition and basal pollen response is the transfer of water from the dry stigmatic papillae cells to that of the desiccated pollen grain. This transfer of water allows for the adhered pollen grain to hydrate, germinate, and grow a pollen tube for the eventual delivery of the sperm cells to the ovule (Preuss et al. 1993; Zuberi and Dickinson 1985). Once hydrated, the pollen tube will emerge at the point of pollen-papilla contact and penetrate the stigma surface between the plasma membrane and the overlaying cell wall (Elleman CJ 1992; Kandasamy MK 1994). Pollen tube entry into the stigma surface acts as a second barrier to select for compatible pollen tubes. After penetration, the compatible pollen tube grows down the base of the stigma, through the style to enter the transmitting tract, and grow between the cells of the ovary to reach the ovules for fertilization. As expected, the pollen-pistil interactions are tightly regulated during growth through the ovary and also in regards to the pollen tube interaction with the ovule (for review see, (Beale and Johnson 2013; Dresselhaus and Franklin-Tong 2013; Higashiyama and Takeuchi 2015).

Previous studies identified, EXO70A1, a subunit of the exocyst, as a key factor involved in the early interaction of the compatible pollen grain and the stigmatic papilla cell. EXO70A1 was determined to be necessary for the acceptance of a compatible pollen grain and *A. thaliana* stigmas in the *exo70a1* mutant display a constitutive Col-0 wild-type pollen rejection phenotype (Safavian et al. 2014; Samuel et al. 2009). This rejection phenotype of the *exo70a1* mutant stigma could be rescued by a fusion construct of Red Fluorescent Protein (RFP):EXO70A1 under a stigma specific promoter, SLR1 (Samuel et al. 2009), or it could be partially rescued in a high relative humidity environment (Safavian et al. 2014). Furthermore, when a stigma-specific RNAi construct was expressed in *Brassica napus* ‘Westar’, acceptance of a compatible pollen grain was impaired (Samuel et al. 2009). These results with EXO70A1 lead to a comprehensive analysis of the other seven of the subunits of the exocyst complex, where stigma specific RNAi silencing constructs were made to knock down the expression of each individual gene for; *SECRECTORY3 (SEC3), SEC5, SEC6, SEC8, SEC10, SEC15* and *EXO84* in *A. thaliana*. This study looked in depth at the impact of each individual subunit through a study of pollen grain adhesion, pollen grain hydration, pollen tube penetration, seed set and overall fertility in these transgenic lines. This exceedingly comprehensive study demonstrated that each of the individual exocyst subunits are necessary for compatible pollen grain acceptance and a fully functional exocyst complex is required for compatible pollen acceptance (Safavian et al. 2015). The results from the exocyst study conclusively demonstrated that polarized secretion via secretory vesicles plays a critical role in pollen grain hydration from the stigmatic papillae cells.

Therefore, to identify other factors that impact secretory trafficking during the process of compatible pollination, we sought to investigate another aspect of the secretory system, retrograde transport. With increased anterograde traffic in the papilla cell to deliver vesicles to the pollen contact point to enable pollen grain hydration, there would be increased retrograde traffic to maintain membrane balance in the system. Coat Protein II (COPII) vesicles are involved in the anterograde transport of protein out from the ER while the Coat protein 1 (COPI) coated vesicles are the mediators of intra-Golgi transport and retrograde transport from the Golgi back towards the endoplasmic reticulum (Ahn et al. 2015; Brandizzi and Barlowe 2013; Gimeno-Ferrer et al. 2017). In regards to the intra-Golgi transport of COPI vesicles the directionality of their transport is still a matter of debate; however, their role in retrograde transport is clear (Jackson et al. 2012). These two types of Coat proteins are able to selectively capture protein cargo from the membrane compartment of origin as well as the fusion machinery necessary for vesicle delivery and formation of the secretory vesicles. It has been shown that COPI vesicles formed at the Golgi help to retrieve ER resident proteins back to the ER. Many of these type I transmembrane proteins have a dilysine-based motif, or variant motifs, that are recognized by the COPI coat (Jackson et al. 2012; Woo et al. 2015). The major component of the COPI coat is the coatomer complex; this consists of seven subunits (α/β/β/γ/δ/ε/ζ) that form two subcomplexes, B-(α/β/ε) and the F-subcomplex (β/δ/γ/ζ), with the B-subcomplex making up the outer layer of the protein coat and the F-subcomplex making up the inner layer (Jackson 2014). The entire complex is recruited to the Golgi membrane *en bloc* and is essential in all eukaryotes (Hara-Kuge et al. 1994). More recent structural studies have emphasized that the subunits are highly connected and the two subcomplexes model does not fit the previously proposed model based on other coat protein structures (Dodonova et al. 2015). What is clear, is that COPI polymerizes on the membrane surface following the recruitment of ADP-ribosylation factor 1 (ARF1), a small GTPase, in its GTP-bound confirmation with uptake of cargo, and that this combination of factors leads to the formation of the COPI vesicles (Jackson 2014). Several studies have demonstrated that the α-COPI and β’-COPI subunits play a role in cargo binding (e.g dilysine motif containing) (Jackson 2014) and that knock-down of ε-COPI resulted in the mislocalization of endomembrane proteins that contain a KXD/E motif and severe morphological changes in the Golgi structure (Woo et al. 2015). In eukaryotes, it has been demonstrated that γ-COPI interacts with ARF1 and that γ-COPI stability requires the presence of ζ-COPI (Jackson 2014).

In plants, the genes that encode the various components of the COPI subunits and machinery have been identified (α/β/β’/γ/δ/ε/ζ) and in *A. thaliana*, several different isoforms of all of the coatomer subunits, except for γ-COPI and δ-COPI have been identified (Ahn et al. 2015; Gao et al. 2014; Robinson et al. 2007; Woo et al. 2015). At the ultrastructural level in *A. thaliana*, two structurally distinct type of COPI vesicles have been identified by electron tomography (Donohoe et al. 2007; Gao et al. 2014). Characterization of the β’-γ-and δ-COPI subunits examined their function in *N. benthamiana* and tobacco BY-2 cells, where it was demonstrated that the COPI complex was involved in cell plate formation (via the phragmoplast), Golgi maintenance and disruption lead to programmed cell death after COPI depletion via inducible gene silencing of *β’-COPI*, *γ*-COPI and *δ*-COPI (Ahn et al. 2015). The α2-COPI isoform was characterized in the context of several α*2-copi* loss-of-function mutants. The *α2-copi* mutants had major defects in overall plant growth and morphology, and the Golgi structure was altered in addition to the subcellular localization of p24δ5, a protein that has been shown to cycle between the ER and the Golgi, which also contains the dilysine motif that is recognized by α-COPI isoforms (Gimeno-Ferrer et al. 2017).

## Materials and Methods

**Plant Materials:** Col-0 wild-type seeds (CS28167) and the COPI T-DNA lines are as follows; At1g62020, α1-COPI (SALK_078465c), At4g31490, β-COPI (SALK_017975 & SALK_084745), At1g52360, β’1-COPI (SALK_206753c), At3g15980, β’2-COPI, (SALK_13730), At1g30630, ε-COPI (SALK_026249) and At4g34450, γ-COPI (SALK_103822). All seed stocks came from the Arabidopsis Research Center (ABRC), Columbus, Ohio, USA.

### Plant growth conditions

All plants were grown under long day conditions, 16 hours light/8 hours dark, with a daytime temperature of 22 °C and a nighttime temperature of 17°C. *A. thaliana* seeds were surface sterilized with 1 volume bleach and 2 volumes Triton X-100 for 15 minutes and then washed with sterile dH_2_O, 4 times. After sterilization all seeds were stratified for a minimum of 3 days at 4°C and were then plated out on ½ Murashige & Skoog, pH 5.8 with 0.4% w/v phytoblend agar plates. Once the seedlings had 2 true leaves, they were transferred to soil (ProMix-BX + mycorrhizal species, Quebec, Canada). The relative humidity of the chambers was recorded to be between 40–70% during all experiments.

**Pollen adhesion, pollen tube growth and seed set analyses:** Pollen grain adhesion and pollen tube growth were observed on freshly opened flowers. To determine the correct age of the freshly opened flowers, stage 12 flower buds (the final bud stage prior to bud opening; (Smyth et al. 1990) from wild-type Col-0 and transgenic plants were emasculated and covered in plastic wrap to prevent drying of the tissues overnight. 24 hours later, individual anthers, from either Col-0 wild-type flowers or transgenic plants, were used to manually apply pollen grains to the stigmas of the emasculated flowers when the papillae were fully elongated. 2 hours-post pollination, whole pistils were removed from the plant and placed in a fixative solution (300µL of ethanol:glacial acetic acid [3:1]) at room temperature for 30 minutes. Fixative was then removed and the pistils were washed with sterile dH_2_O three times and then incubated in 300µL of 1N NaOH for 1hr at 60 °C. Next, the 1N NaOH was removed and the pistils were washed with sterile dH_2_O three times and then stained with 300µL of 0.1% (w/v) aniline blue at 4°C overnight. To visualize pollen grain adherence and pollen tube penetration, whole pistils were mounted on slides with mounting media (4% (w/v) propyl gallate) to prevent photobleaching. Images were captured with a Zeiss Axio-observer fluorescence microscope. At minimum, 10 pistils were imaged per each individual line for pollen grain adherence (Brightfield) and pollen tube growth (DAPI-358/463nm). Reciprocal crosses were conducted by pollinating wild-type Col-0 pistils with pollen from the copi mutant transgenic lines to confirm the pollen viability of the transgenic lines. The pollination assays were performed at a relative humidity level of between 45–65% and were monitored with a digital hygrometer in the chamber at all times. To determine seed set, the number of seeds per silique were tallied for Col-0 and each independent transgenic line (n=30). Seed set treatments were examined using Welch’s ANOVA after running Levene’s test for homogeneity. For comparisons between genotypes, we used Dunnett’s multiple comparisons test and a Tukey post-hoc test using the SAS for University software package with an α =0.05 for a treatment (in this case genotype) to be determined as statistically significant (Cary, NC).

**Genotyping of T-DNA lines and RT-PCR analysis:** The *A. thaliana* T-DNA insertion mutants, were genotyped by PCR (primers in Supplemental Table 1). Homozygous individuals were identified for all lines expert for SALK_103822. SALK_103822 individuals were heterozygous mutants and these results indicate that this mutant has a homozygous lethal phenotype. To confirm that the T-DNA insertion lines were null mutants, RNA was extracted from pistils from each line using the total Plant/Fungal RNA extraction kit (Norgen, Niagara, ON) and then quantified using a Nanodrop spectrophotometer (GE Nanovue Plus). 1µg of total RNA was treated with DNase to remove any potential DNA contamination for 15 minutes with DNase I (Fermentas) followed by 25mM ETDA at 65 °C. 500 ng total RNA was then used as input in cDNA synthesis with SuperScript III plus RNaseOUT (Invitrogen). Primers for the RT-PCR analyses are listed in supplemental table 1. All PCRs were performed in a ABI 9600 thermocycler with the following program; step 1 - 94°C for 5 minutes; step 2 - 94°C – 30 sec, primer Tm 54–59°C – 30 sec - 72°C – 45 seconds for 25–28 cycles, step 3 – extension 72°C – 10 minutes, step 4 – hold at 10°C. Taq polymerase (BioBasic, Amhyrst, NY) was used with the manufacturer’s instructions for all PCR analyses.

## Results

### Genotyping of *copi* mutant *A. thaliana*

To characterize the potential role of the COPI complex in the stigimatic papillae following compatible pollination, we acquired a wide variety of *copi* T-DNA mutants in different subunits. In summation, we characterized seven different T-DNA lines in five different COPI subunits and each one is shown in Figure 1. The presence of the T-DNA was confirmed by genotyping of each individual (see Methods). We examined α1-COPI (At1g62020) in the SALK_078465c line (Fig 1A), which has the T-DNA inserted in the third exon of the coding sequence and we were able to isolate homozygous mutant plants in α*1-copi*. For β-COPI (At4g31490) two homozygous mutant SALK T-DNA lines (Fig 1B) were isolated and genotyped. For clarity, they will be referred to as follows, β*-copi-1* (SALK_017975) and β*-copi-2* (SALK_084745) alleles. Both alleles had the T-DNA insertion in the second exon of the coding region of the gene. For β’-COPI two different paralogues were examined (Fig 1C), β’*-1-copi (*At1g52360) and β’*-2-copi* (At3g15980)*. β’-1-copi* had a T-DNA insertion in the fifth exon (SALK_206753) and *β’-2-copi* had the T-DNA insertion in the second exon (SALK_135730). *ε-COPI* mutant (At1g30630) had a single T-DNA in the 5’UTR (SALK_026249) as shown in Fig 1D. Lastly, the γ-COPI (At4g34450) was examined for the T-DNA in the eighth exon (SALK_103822) as shown in Fig 1E. Genotyping of the *γ-copi* mutant plants revealed that only heterozygous mutants were recovered, indicating that the homozygous *γ-copi* mutant is likely lethal. Therefore, all of the characterizations were performed on heterozygous individuals that were always genotyped before experiments to confirm that they were indeed heterozygous and not wild-type.

**Fig. 1.**
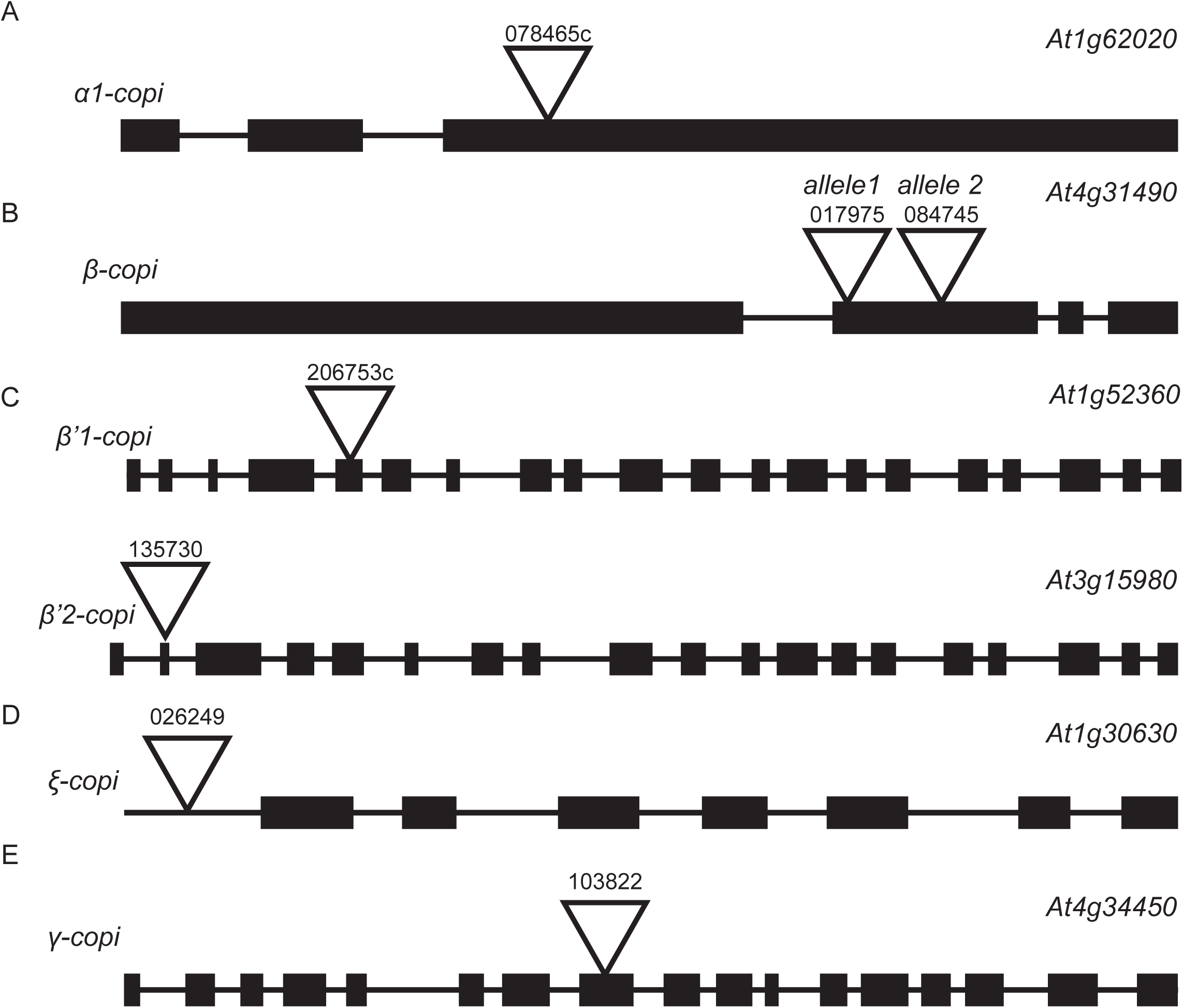
COPI T-DNA lines used in this study. All constructs are SALK T-DNA lines that were genotyped to determine the presence of the T-DNA insert and to determine if the individual was homozyogous for the T-DNA. All of the T-DNA insertion points are indicated by the triangles. All mutants had insertions in exons except for the ε1-copi mutant where it was present in the 5’ region of the gene. A.) *α1-copi* mutant in At1g62020 with SALK line 078465c. B.) Two different T-DNA insertions were examined for *β-copi*, SALK 017975 & 084745 both in the second exon. *C.) β’-copi* mutants were in two different copies of β-COPI, At1g52360, SALK 206753c and At3g15980, SALK 135730. D.) *ε-copi* mutant of At1g30630 with the T-DNA line of SALK 026249. E.) *γ-copi* mutant in At4g34450 with the SALK line 103822. Interestingly, in SALK line 103822, we could only recover heterozygous indicating that the line is homozygous lethal

### RT-PCR confirmation of knock-out or knock-down of *COPI* genes

After genotyping the *copi* T-DNA plants, each line was examined to verify the knock-out or knock-down of each individual *COPI* gene (Fig 2). As the major focus of this study is on the role of the COPI complex in the pistil, pistils were collected from T2 siblings for each line and pooled to have enough tissue for subsequent analyses. Before examining the expression of each *COPI* gene individually, Col-0 wild-type cDNA was compared with that of the transgenic lines generated with the reverse transcriptase (RT+) and without reverse transcriptase (RT−) to confirm that the DNase treatment was successful and there was no genomic DNA contamination was found in the samples. Figure 2A shows the expression of *Elf1α*, the housekeeping gene that was selected for as the internal control analyzed at 25 cycles. *Elf1α* showed equal levels of (non-quantitative) expression across all samples, and no genomic contamination in the samples as demonstrated by no signal in the RT-samples. Gene specific primers were designed to the 3’ end of each *COPI* gene and were made to flank an intron so that genomic contamination would be easy to detect. Then the expression of each gene was compared between Col-0 and the mutant lines and all samples were examined at 25 cycles, the same as the *Elf1α* (Figure 2B). The *copi* mutant lines were true knock-outs in the following subunit isoforms; *α1-COPI*, *β-COPI*, and *ε-COPI*. Interestingly, neither *β’1-COPI* nor *β’2-COPI*, were knock-outs of gene expression. Even though *β’1-COPI* and *β’2-COPI* were not knock-outs, the lines were characterized to determine if they would show a phenotype that was milder than the other *copi* mutants or if the lines appeared wild-type. As the *γ-copi* appeared to be homozygous lethal during the genotyping of the T-DNA plants, we characterized the heterozygous individuals (genotyping each individual) and when examining the expression of *γ-COPI* with RT-PCR, it was not knocked down as expected. As our studies continued, we did observe phenotypes which were a result of the T-DNA mutation, even though they were heterozygous and not homozygous knock-outs.

**Fig. 2.**
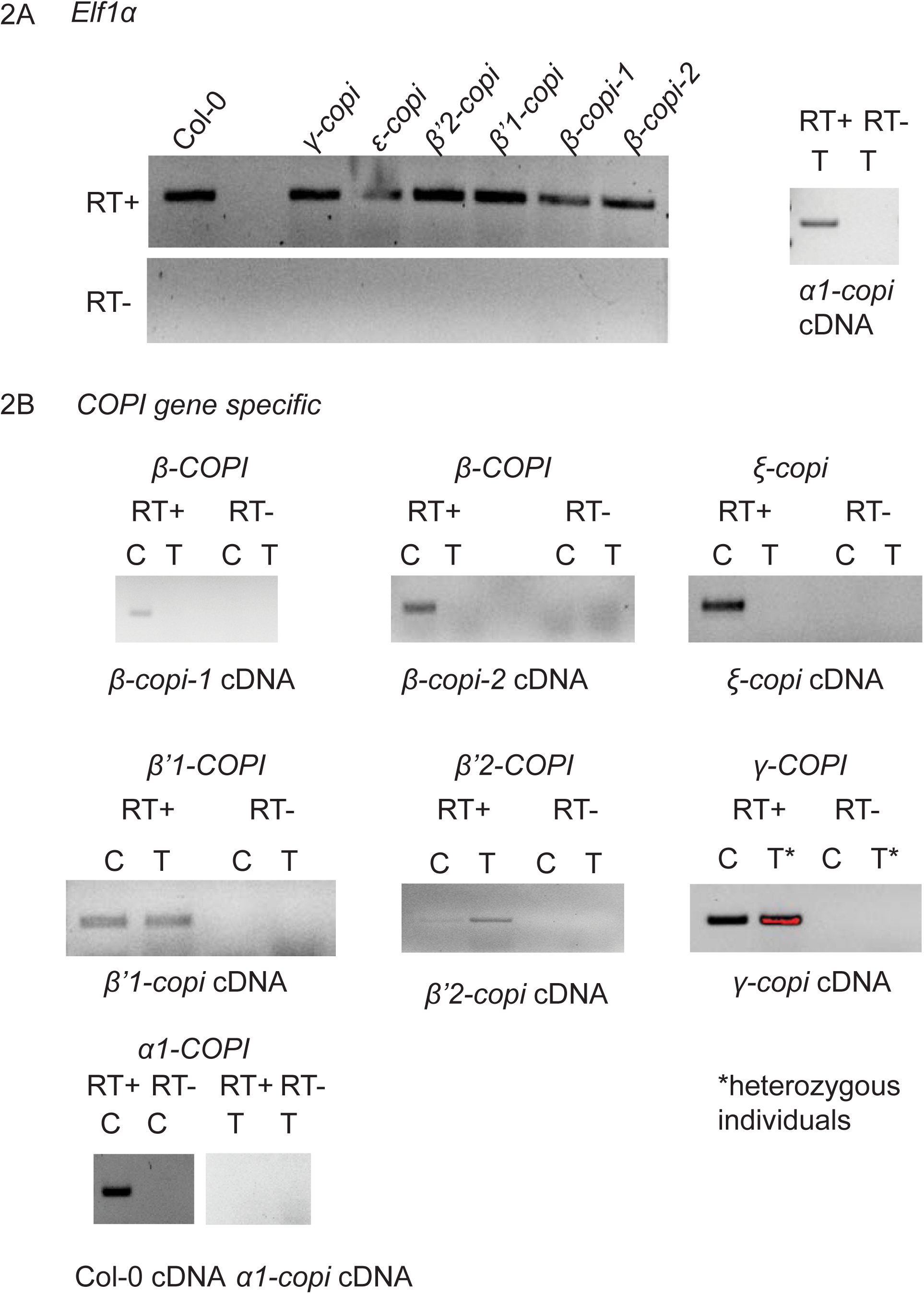
Verification of COPI isoform gene expression to detect knock-out or knock down by the T-DNA insertion. 2A.) *Elf1a* as a control for housekeeping gene expression levels and verification of the RT-samples to confirm there was no gDNA present in samples. 2B.) RT-PCR to examine the knock-out or knock down of each individual *COPI* isoform. The gene analyzed is shown above the image and the cDNA input below. C= Col-0 cDNA, T= T-DNA cDNA, * notes transgenic individuals that were determined to be heterozygous for the T-DNA

### *copi* mutant stigmas show a wide range of compatible pollen response phenotypes

It has been previously demonstrated that compatible pollinations are facilitated by directed secretory activity to the point of pollen contact with the individual stigmatic papilla cell (Doucet et al. 2016; Elleman and Dickinson 1996; Goring 2018; Indriolo and Goring 2014; Indriolo et al. 2014; Safavian and Goring 2013; Safavian et al. 2015; Samuel et al. 2009). At 10 minutes post-pollination in *Arabidopsis spp*., TEM images clearly show that secretory vesicles fuse with the plasma membrane at the point of pollen contact (Safavian and Goring 2013) and that when incompatible pollen grains are rejected, this process is driven by the removal of secretory vesicles at the pollen contact point through autophagy (Indriolo et al. 2014; Safavian et al. 2015). The requirement for the exocyst complex to deliver secretory vesicles to the pollen contact point, was clearly demonstrated a through and robust examination of exocyst knock-out mutants and RNAi mediated known-down in *A. thaliana*. As the exocyst derived vesicles are a result of anterograde vesicle action, it is clear that increased secretory activity is required for compatible pollen acceptance (Doucet et al. 2016; Goring 2018; Safavian and Goring 2013; Safavian et al. 2015). With increased secretory activity towards the pollen contact point, there would need to be an equal amount of secretory activity away from the pollen contact point to maintain membrane balance within the endomembrane system (Goring 2018).

Therefore, to determine if each individual subunit and isoform of the COPI complex analyzed in this study is necessary for pollen grain hydration, manual pollinations followed by Aniline Blue staining were performed 2 hours post-pollination. All floral buds were emasculated at stage 12 (Smyth et al. 1990) and covered with plastic wrap to prevent dehydration and prevent premature pollination while in the growth chamber. The following morning, *copi* mutant stigmas were gently manually pollinated with Col-0 pollen, and the relative humidity was recorded at the time of pollination. All results presented here were performed at a relative humidity that was higher than desired, ranging between 59–71% and most of the time between 62–64%. It was previously determined that a relative humidity of less than or equal to 40% prevents spurious pollen grain hydration from the ambient humidity as opposed to from the stigmatic papillae cells (Safavian et al. 2014). 2 hours after pollination, entire pistils were collected and stained with Aniline Blue to determine pollen grain adherence and pollen tube penetration into the stigma and style. All pollinations were consistantly compared to the control pollination of a Col-0 wild-type stigma pollinated with Col-0 pollen (Fig 3A & E). Wild-type pollinations show hundreds of pollen grains that have adhered to the stigmatic papillae cells in the brightfield image while the Aniline Blue stain shows the germinating pollen tubes growing towards the ovules in the ovary (not visible in images). The COPI subunit with the strongest and most striking phenotype is that of the *α1-copi* (Fig 3B & F), which clearly shows a low number of pollen grains on the stigma surface. The few pollen tubes that have germinated are very short and have not grown very far when compared to that of the Col-0 control pollen tubes. Additional Aniline Blue and Brightfield images of the *α1-copi* stigmas (Supplemental Fig 1) show low pollen grain adherence and either short delayed pollen tube growth or almost none at all. The second strongest phenotype is the one observed in the *γ-copi* line (Fig 3L & P), which interestingly does not seem to have the same pollen grain attachment phenotype as the *α1-copi*, with considerably more pollen grains adhering to the stigma but with almost no pollen tube germination or growth at 2 hours post-pollination, again compared to a Col-0 wild-type pollination. The *γ-copi* line was consistent in its phenotype with additional images showing poor pollen tube growth and germination with the Aniline Blue stain (Supplemental Fig 1) but pollen grain adherence similar to Col-0 wild-type. It is quite clear that the *α1-copi* and the *γ-copi* stigmas have a noticeable defect in compatible pollen grain acceptance, pollen grain germination, and pollen tube penetration. A milder phenotype was observed in the *ε-copi* stigmas (Fig 3K & O & Supplemental Fig 1). Pollen grain attachment appears wild-type, as shown in the brightfield image and the pollen tube growth appears weaker with shorter tubes and less density in the fluorescence signal of the stained tubes. In contrast to the *α1-copi*, *γ-copi*, and *ε-copi* stigmas, the *β-copi* (2 alleles) and *β’-copi* (2 genes) mutant stigmas appeared as though they had the same phenotype as Col-0 stigmas (Fig 3 C, D, G-J, M & N & Supplemental Fig 1). The *β-copi* and *β-copi* stigmas had wild-type like pollen grain attachment. The pollen tube germination and growth were robust and rapid similar to Col-0 wild-type pollen. Therefore, the Aniline Blue stains show a gradient of phenotypes in pollen grain adherence and pollen grain germination between the different mutants of COPI subunits early in pollen grain recognition and acceptance. This may be in part explained by the fact that certain lines are not true knock-outs in the expression of the gene e.g. *β’2-copi* or the fact that these experiments were conducted at a higher than desirable relative level of humidity (60–70%) (Safavian et al. 2014).

**Fig. 3.**
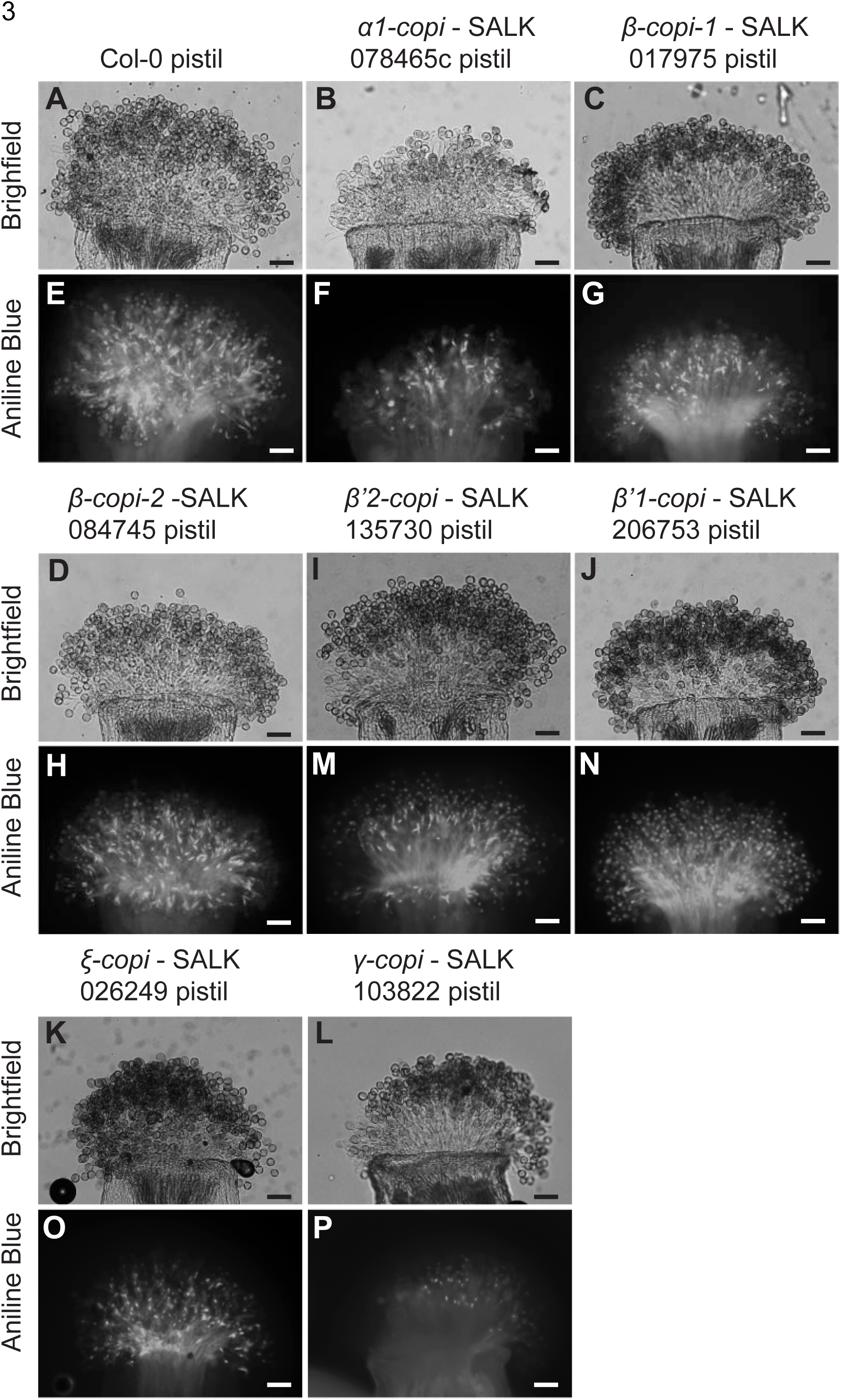
Pollen grain attachment and pollen tube growth following manual pollination of pistils from COPI subunit knock-out lines with wild-type Col-0 pollen. A & E, wild-type Col-0 pollen on a Col-0 pistil. B-D & F-P show, Col-0 pollen on *copi* mutant stigmas at 2 hours post-pollination. Images A to P represent the average observed phenotype for each line examined. Each line was tested three times and the additional two panels display a similar phenotype (Supplemental Figure 1). Two subunits, *α1-copi* (B & F) and *γ-copi* (L & P), show drastic reduction in pollen grain adherence and pollen tube germination and growth, when compared to Col-0 wild-type pistils. In contrast, most of the lines show close to wild-type or wild-type pollen grain adherence and pollen tube penetration. The *ε-copi* (K & O) line shows normal pollen grain adherence and retarded pollen tube penetration into the pistil with shorter pollen tubes when compared to the other more robust lines. Scale bars = 50 μm

### Certain *copi* mutant pollen lines have difficulty germinating on stigmas

Working with T-DNA knock-out lines of the COPI complex allowed for the phenotype of the pollen grains to be examined in the context of a compatible pollination on wild-type Col-0 stigmas. This allowed us to tease out the role of the COPI complex in pollen-pistil interactions extending to the pollen grain and the pollen tube a metabolically active structure undergoing high rates of secretory activity driving tip growth (Chebli et al. 2012; Cheung and Wu 2008; Luo et al. 2017). All of the *copi* mutant pollen grains were compared to that of wild-type Col-0 pollen on a Col-0 stigma (Fig 4A & E) and Col-0 stigmas were pollinated by *copi* mutant pollen (Fig 4 B-D, F-P). Overall, there was not a very clear difference in pollen grain adherence with the *copi* mutant pollen for most of the subunits when examined visually under brightfield. Additionally, the only phenotypes that were observed were at most mild; this includes the *α1-copi* pollen, with pollen tubes that appeared to be shorter than all of the other pollen tubes when examined for Aniline Blue stain (Fig 4B & F), and *β-copi-2* (4D& H) and *γ-copi* (4L & P). For the *α1-copi*, *β-copi-2*, and *γ-copi* pollen on wild-type stigmas the shorter pollen tube phenotype was visible in all images surveyed (Supplemental Fig 2). An interesting observation is that of the two different T-DNA knock-out alleles that were examined, only one of the alleles showed a pollen specific phenotype; *β-copi-2* (SALK_084745) while *β-copi-1* (SALK_017975) appeared similar to Col-0 wild-type. Both T-DNA knock-outs were confirmed by the RT-PCR analysis (Fig 2) which rules out an obvious difference in the degree of knock-out between the two different T-DNA lines. Both of the T-DNA insertions were also confirmed to be in the second exon and are relatively close to one another. Further analysis will be needed to determine what effects the insertion difference may have on the impact of the mutation on germinating pollen grains during compatible pollinations. All of the other *copi* mutant lines appeared indistinguishable from that of the wild-type Col-0 pollen. As with the examination of the *copi* mutant stigmas, the relative humidity of the growth chambers during these experiments was between 60–70% and it would have been preferable to have a lower level of humidity; however, both the age of the growth chambers, and the evaporative cooler that is used to regulate the temperature of the room, contributes to a higher level of relative humidity than with refrigerated air (Safavian and Goring 2013; Safavian et al. 2015).

**Fig. 4.**
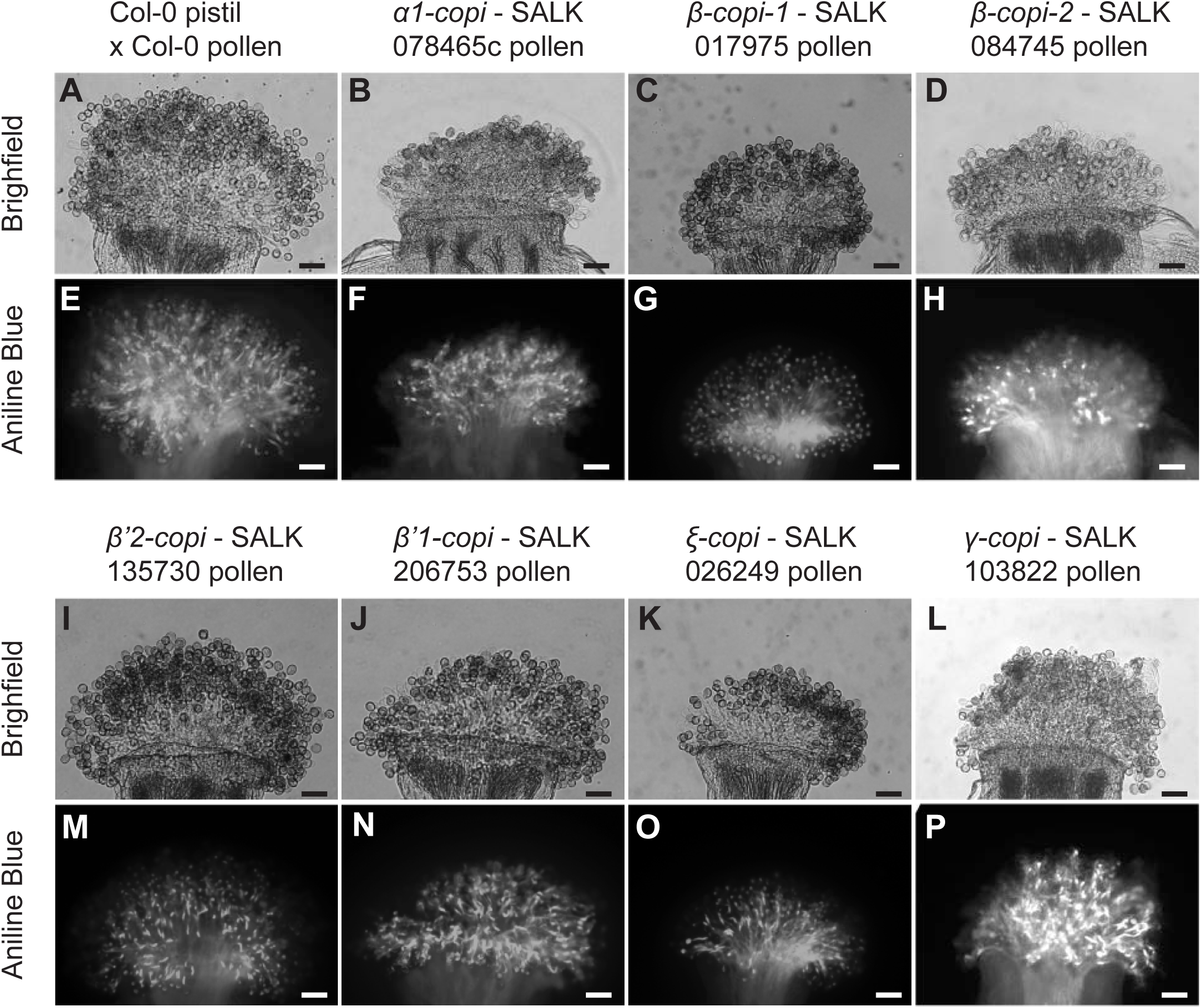
Pollen grain attachment and pollen tube growth following manual pollination of pistils from wild-type Col-0 with COPI subunit knock-out line pollen. A & E, shows the control Col-0 pollen on Col-0 stigmas at 2 hours post-pollination. Images B-D, F-P show Col-0 pistils that have been manually pollinated with copi knock-out line pollen grains, 2 hours post-pollination. All images represent the average observed phenotype for each line examined. Each line was tested three times and the additional two panels display a similar phenotype (Supplemental Figure 2). Several of the *copi* knock-out lines appear to have delayed or reduced pollen grain adherence and pollen tube penetration including *α1-copi* (B & F) *β-copi* (D & H), *ε-copi* (K & O) and *γ-copi* (L & P). Scale bars = 50 μm

### Seed set in *copi* mutant lines can range from severe to wild-type depending on the subunit examined

Aniline Blue stains are an excellent way to access the early events during pollen acceptance and pollen grain germination on mutant stigmas. The stains capture the pollen grain’s interaction with the stigmatic papillae cells within the first 2 hours of the flower being receptive to pollen. However, this does not give insight into later events within this process, such as the successful fertilization of the female gametophyte by the sperm cells. To examine these later events, stage 12 floral buds (Smyth et al. 1990) were emasculated and the following morning were manually pollinated with Col-0 pollen (1–2 anthers per stigma). The plants were then returned to the growth chamber and allowed to develop for 10 to 11 days. As with the other experiments, the relative humidity of the growth chambers was between 60–70% during the time period. At day 10 or 11 post pollination, the siliques were removed from the plants and were dissected to determine the seed set. Seed set was tallied for 30 siliques for each individual *copi* mutant line and compared to Col-0 self-pollination under the exact same environmental conditions (Fig 5). Col-0 pollinations had an average seed set of 53.2 (+/−3) seeds per silique and these served as our control for the rest of the pollinations. The *copi* mutant line with the most severe reduction in seed set was that of *α1-copi*, as this line had an average seed set of 3.0 (+/-6.1) seeds per silique. These results indicate that the knock-out of *α1-copi* not only has difficulty in short term pollen-pistil interactions, but that this continues as development progresses and Col-0 wild-type pollen cannot overcome the deficiencies of the *α1-copi* pistil. Another line, showing a decrease in the number of seeds per silique was that of the *γ-copi* line with 37.3 (+/−7.9) seeds per silique. But when compared to Col-0 wild-type and the *α1-copi* line, the *γ-copi* plants are much more similar to that of Col-0 wild-type pistils than the *α1-copi* phenotype. All remaining lines showed seed set numbers that were very close to wild-type or indistinguishable from that of wild-type pollinations. The phenotypes are presented in order of increasing amounts of seed set; *ε-copi* had 41.6 (+/−4.2) seeds per silique, *β-copi-1* had an average seed set of 42.4 (+/−12.1), *β’2-copi* showed 46.4 (+/−15.2), *β’1-copi* had an average of 49.9 (+/−6.1) seeds per silique and lastly, the *β-copi-2* siliques, had on average 57.1 (+/−4.7) seeds per silique. Despite examining a total of 30 siliques per each transgenic line, the *β-COPI*, *β’-COPI* and *ε-COPI* knock-outs had unequal variance was accounted for during the analysis of the seed set data. Therefore, seed set treatments were examined using Welch’s ANOVA after running Levene’s test for homogeneity. For comparisons between genotypes, we used Dunnett’s multiple comparisons test and a Tukey post-hoc test. The results of the statistical analyses are as follows; *α1-copi*, was statistically significantly different than Col-0 wild-type and all of the other mutant lines (α=0.05 with Tukey adjustment). Therefore, the most severe phenotype observed of all of the COPI mutant subunits in regards to seed set was that of the *α1-copi* in line with the Aniline Blue stain results for early pollination phenotypes. The *γ-copi* was significantly different from Col-0 and all of the other mutant lines except for the *ε-copi*, which was similar to that of the *γ-copi*. Therefore, the phenotypes of the *γ-copi* and *ε-copi* mutant lines were less severe than that of *α1-copi* when analyzed by seed set but were still different from Col-0 wild-type. The *ε-copi* was more subtle than *γ-copi* as it was similar to *β-copi-1*. Not surprisingly, Col-0 was similar to *β-copi-2*, and both of the mutant isoforms of *β’-copi*. In conclusion, the seed set experiment has allowed for the resolution of much more subtle differences in the response of certain COPI isoforms in the stigma when pollinated with Col-0 wild-type pollen.

**Fig. 5.**
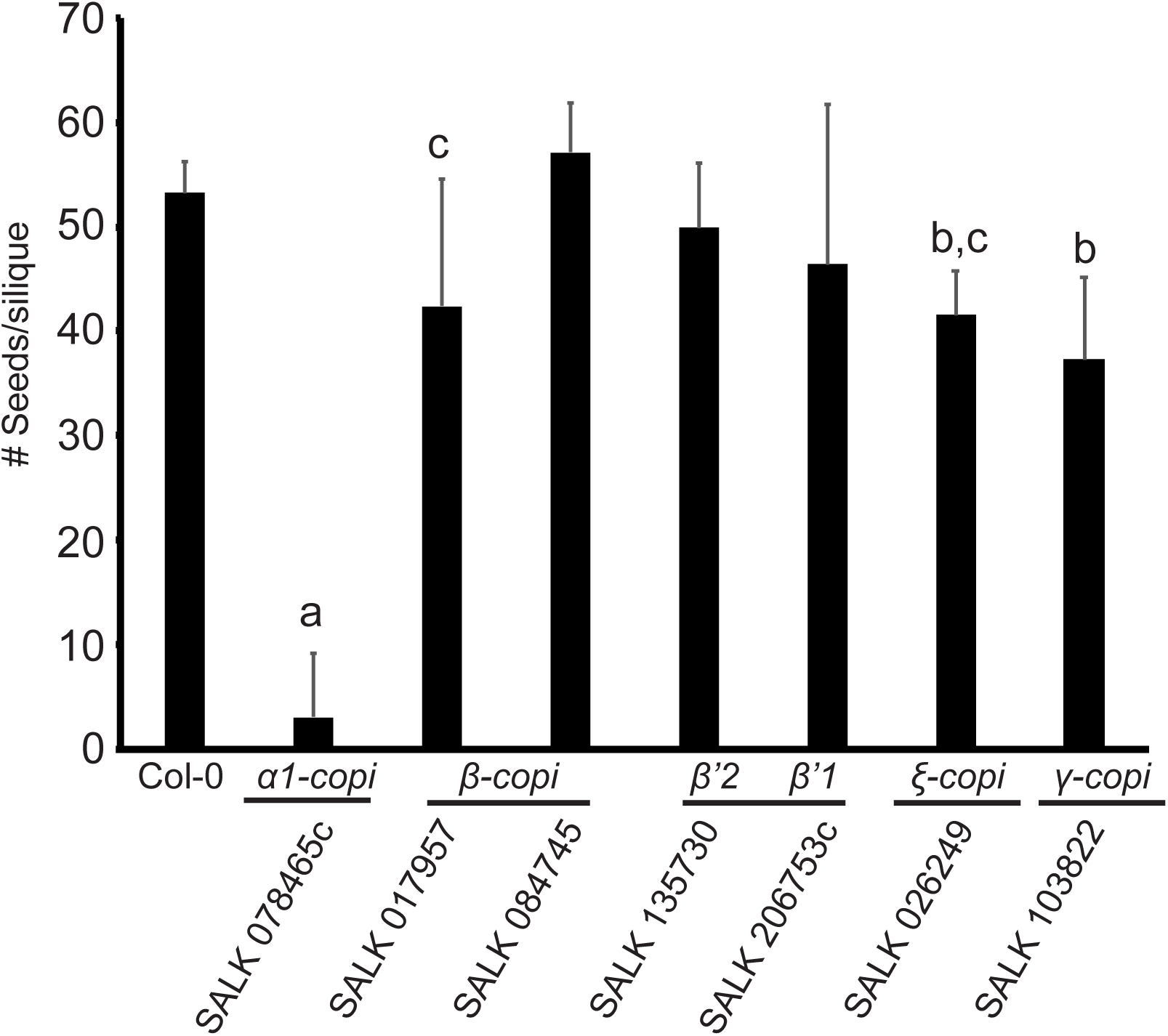
Graph showing the number of seeds per silique for Col-0 and each *copi* mutant line. All pistils were pollinated with wild-type Col-0 pollen (n=30 for each plant line). The mean number of seeds per silique for average silique sizes in each plant line, following manual pollination. All seeds were tallied 10 to 11 days after manual pollination. Error bars = SE. A., indicates that the *α1-copi* line is statistically significantly different from the Col-0 wild-type plants and the rest of the *copi* T-DNA mutant lines. This was examined with a one-way ANOVA with no significant difference between blocks but a significant different between genotypes (*α* = 0.05) followed by a Dunnett’s and Welch’s test with Tukey post-hoc test. B. indicates lines that are statistically similar to one another but different from all remaining lines and C. indicates lines that are statistically similar and different than all other lines

## Discussion

This is the first study examining the role of the COPI complex in the context of pollen-pistil interactions. Five individual subunits, two isoforms of *β’1-COPI* and *β’2-COPI* and two different mutant alleles of a *β-COPI*, that comprise the COPI complex, were analyzed for their role in the pistil of *A. thaliana* during compatible pollination and pollen germination on the stigma surface. The subunit with the strongest phenotype when knocked out by the presence of a T-DNA mutation was *α1-copi*. Through all of these studies, *α1-copi* displayed a clear reduction in pollen grain adherence, pollen grain germination and pollen tube growth when Col-0 wild-type pollen grains were applied to the *α1-copi* stigmas. These results demonstrated that during early events, defined as the first 2 hours following a compatible pollination, can be disrupted by a mutation in a COPI subunit. Furthermore, when looking events that occur much later than the early events, the *α1-copi* plants continue to have issues that negatively impacted the interaction of the pollen tube with the pistil. This is shown through the seed set experiments that used the manual pollination of the *α1-copi* stage 13 flowers with Col-0 pollen, which were allowed to progress 10–11 days before dissecting the siliques to tally the number of viable seeds that had developed. Any early issues among the pollen grain’s interactions with the mutant stigmas should have been overcome within this longer time frame, yet overall, the average number of seeds per silique was 3.0 (+/−6.1) and was demonstrated to be statistically significant.

Even though the initial goal of our experiments was to examine the potential impact of a COPI mutation from the pistil side of a compatible pollination, the impact of the mutation was examined in the mutant pollen grains that were applied to Col-0 wild-type stigmas. The *α1-copi* mutant pollen showed issues with pollen grain adherence, hydration, germination and pollen tube growth within the first 2 hours following the manual pollination. It is quite clear that a mutation in *α1-copi* has a strong impact on the haploid male gametophyte, as well as the diploid female stigma, style and ovary. These phenotypes were likely discovered due to our focus on the role of this isoform *α1-copi* in the context of pollen-pistil interactions while previous characterizations of this line were reported to appear wild-type when grown vegetatively. Previously, when the *α1-copi* plants were compared to the *α2-copi* isoform loss of function mutants, the *α1-copi* plant did not show the *α2-copi* isoform’s phenotype which had global defects in plant growth, resulting in smaller rosettes, stems and roots as well as other morphological problems in protein localization and the Golgi structure (Gimeno-Ferrer et al. 2017). Under our growing conditions, the *α1-copi* plants appeared healthy and wild-type in both size and in overall morphology; these observations are consistent with previous studies of the same mutant line when *α1-copi* plants were compared to the *α2-copi-1, α2-copi-2 and α2-copi-3* plants that showed the dwarf phenotype (Gimeno-Ferrer et al. 2017). As the dwarf phenotypes and the impact on secretory pathway genes in the *α2-copi* plants would alter all aspects of plant growth, future experiments should focus on knocking down *α2-COPI* under as stigma specific promoter such as SLR1 driving a *α2-COPI* RNAi construct in the *α1-copi* background (Trick 1990). This experiment would allow for one to determine if the *α1-copi* phenotype during pollination is, in part, being compensated by the presence of a wild-type isoform of α2*-*COPI.

A subunit that displayed a more subdued phenotype was that of the *γ-copi* mutant; recall that when this line was homozygous, the mutant was lethal under our growing conditions and only heterozygous individuals were characterized. The heterozygous plants were characterized as previous studies showed that VIGS induced gene silencing of the *N. benthamiana γ-COPI* homologue resulted in acute plant cell death, partially due to disassembly of the Golgi apparatus, so it was likely that the *γ-copi* mutant would have a pollen-pistil interaction phenotype (Ahn et al. 2015). As demonstrated by the RT-PCR results, expression of *γ-COPI* was similar to Col-0 wild-type; however, the heterozygous plants displayed a pollination phenotype. This was clearly shown with the Aniline Blue stains as the early events that regulate the adherence and hydration of pollen grains are disrupted by a failure or delay on the papillar side of this interaction. Our experiments were unable to determine how the *γ-copi* heterozygous mutants were able to somewhat overcome the longer term pollination phenotypic effects that were observed with the *α1-copi*. Statistically, the *γ-copi* mutant stigmas were different than Col-0 wild-type and the *β-copi* and *β’-copi* lines with a less drastic reduction in seed set compared to *α1-copi*. Previous studies have shown that γ-COPI directly interacts with ARF1 and with one functional copy of γ-COPI present, it may be able to slowly overcome the lower amounts of protein found due to the mutation.

A factor that may have contributed to the seed set results could be the relatively high humidity found in the growth chambers (between ~50–70%); it has previously demonstrated that higher levels of relative humidity can mask pollen rejection phenotypes in the instance of self-incompatibility or provide enough moisture to overcome an insufficient amount of water provided to the pollen grain from a mutant stigma (Safavian and Goring 2013; Safavian et al. 2015). The facility where all of the plants from these studies were grown in did not have refrigerated air conditioning, which removes humidity from the room, and instead the facility is cooled by an evaporative cooler (daytime highs outside may reach as high as 42 °C and the growth chamber room was at 27 °C and the chambers maintained a constant temperature of 22°C during the day). The evaporative cooler kept the relative humidity in the area of the growth chambers between 58–71% and by design it was not possible to remove this excess humidity from the building. The growth chambers were unable to compensate for the high humidity of the room and showed a similar level of relative humidity throughout this time period.

Another likely factor that may have contributed to the weaker phenotype may be due to the inability to work with a true homozygous knock-outs. Therefore, as stated previously, the next step would be to develop a RNAi construct that is stigma specific (SLR1 driven) to *γ-copi* to transform into the current heterozygous background and to characterize it during stage 13 flowers when the stigmatic papillae cells are receptive to pollen grains.

Analogous to that of the *γ-copi*, similar issues with the high relative humidity of the growth chambers, and room where they were located, likely impacted the studies of ε-COPI, where there were early pollen-pistil phenotypes that were overcome by time and resulted in normal seed set. *ε-copi* pistils showed slightly retarded pollen tube growth when pollinated with Col-0 wild-type pollen and the *ε-copi* pistils had significantly different seed set compared to wild-type. Lastly, the *ε-copi* pollen grains had delayed pollen tube growth and penetration when examined for early pollen-pistil interactions.

Upon examination of the two knock-out alleles of *β-copi*, SALK lines 017975 and 084745, we observed no phenotypes related to the stigma during pollen-pistil interactions. Pollen grains adhered to the stigmatic papillae with no issues, and pollen grain germination and tube growth was robust. The only difference observed was with the *β-copi-2* allele having a mild pollen tube growth retardation in the mutant pollen on wild-type stigmas. Furthermore, the seed set in the *β-copi-2* was statistically similar to that of the Col-0 wild-type pollen while the *β-copi-1* was different then Col-0 wild-type, *β-copi-2* and the *β’-copi* lines, although it was similar to that of the *ε-copi* seed set. When examining the *A. thaliana* genome sequence, another copy of *β-COPI* was identified, right next to this isoform. Based on its location in the *A. thaliana* genome, it appears that these copies of *β-COPI* are the result of a very recent tandem duplication. The sequence of At4g31490 (*β-copi* knock-out lines), when compared to that of At4g31480, the coding sequence is 96.52% similar at the nucleotide level between these two genes. It is highly likely that with such high a level of similarity between these two genes, they code for almost identical isoforms of β-COPI, and the second gene is able to compensate for the knock out of the other. Furthermore, there is precedent for this possibility as when various subunits of the exocyst complex were characterized in compatible pollinations of *A. thaliana*, phenotypes were only observed when a knock-out mutation(s) were crossed with each other or the addition of a RNAi construct was added to knock down the expression of another copy of the same gene (Safavian et al. 2015). Therefore, the next logical step in the characterization of *β-COPI*, and its role in pollen-pistil interactions will require the use of higher order knock-outs. Interestingly, the subtle pollen phenotype for one of the knock-out alleles, *β-copi-2*, may be a result of higher expression of this isoform in the pollen grain and it might play a role in the male gametophyte, which would not be observed in the diploid pistil.

The characterization of the *β’-COPI*, isoforms were not informative, as both of the lines that were selected did not have a knock-out in the expression of these genes. Even though both genes, At1g52360 and At3g15980, were homozygous for the T-DNA insertions in exons, when analyzed by RT-PCR, expression of the genes was either similar to that of Col-0 wild-type, as shown in At3g15980, or had an even higher level of expression than that of the wild-type in At1g52360. Thus, it is not surprising that both of these knock-out lines are not true knock-outs and, when examined by the Aniline Blue stains and seed set assays, they behaved as though they were similar to wild-type plants, with normal pollen grain adherence, pollen grain germination, pollen tube growth and seed set. To characterize these isoforms, new mutant lines will have to be selected and likely combined with the use of RNAi construct to target additional isoforms in a knock-out mutant background.

The results presented here are the first study to examine the role of the COPI complex during compatible pollinations. In the instance of the α-COPI isoforms, we were able to demonstrate the role of the α1-COPI isoform during compatible pollinations in the stigma and that it may have a different role compared to that of the previously characterized a2-COPI isoform, which appears to play a much greater role in secretory traffic throughout development in *A. thaliana* (Gimeno-Ferrer et al. 2017). Many of the other subunits, and their isoforms, were much less informative with more subtle phenotypes during the characterization of their roles during pollen-pistil interactions. These subtle phenotypes are likely due to the fact that other isoforms or additional copies of the genes were able to compensate for the loss of a single isoform (Safavian et al. 2015). As secretory activity and trafficking are essential for regular cellular functions, it will require the use of more precise methods such as tissue specific or inducible promotors to knock down COPI activity in the background of higher order knock-out mutants in *A. thaliana*.

Funding: Support for the Indriolo lab was provided by university start-up funds provided to Emily Indriolo from NMSU.

Support for undergraduate researcher M. Isabella Chavez was provided by the Howard Hughes Medical Institute, Research Scholars Program: #52006932 and #52008103

## Acknowledgements

DACG, MIC and EI designed and performed all of the experiments. EI and DACG performed the statistical analysis. EI wrote the manuscript; DACG and MIC critically edited the manuscript. DACG, MIC and EI contributed to the design of all of the figures and EI performed the final edits of the figures. All authors declare that they have no financial or personal conflicts regarding the results presented in this manuscript. Funding to support the research came from NMSU Start-up funds to EI and undergraduate research scholar funds awarded to MIC through the HHMI research scholars program (#52006932 and #52008103). The authors would like to than the input of Michael Balogh for critical review of the manuscript.

**Supplemental Table 1.**
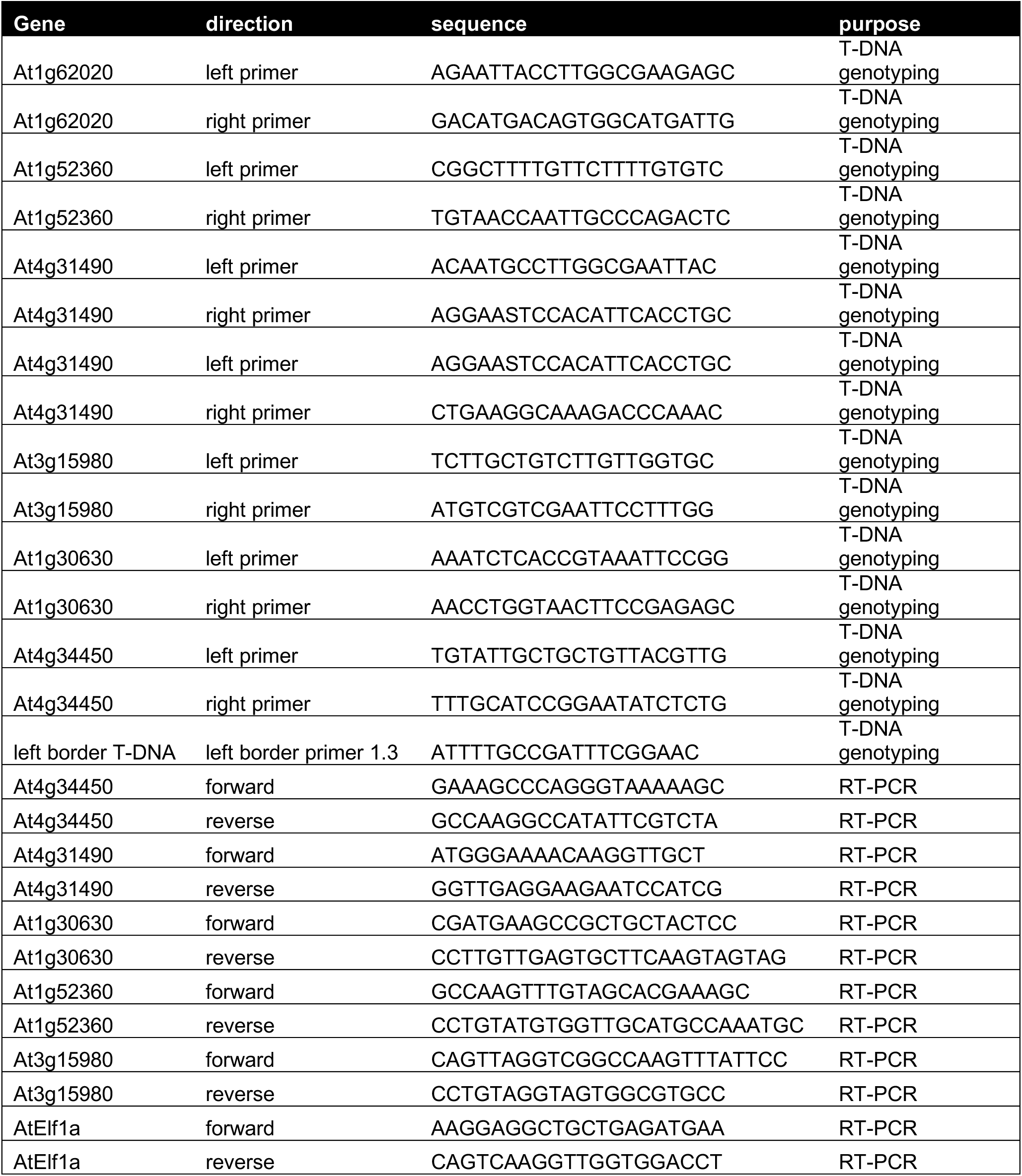
List of primers used in this study. The gene detected by each primer is indicated as well as if the primers were designed for genotyping or for RT-PCR

**Supplemental Fig. 1.** Pollen grain attachment and pollen tube growth following pollination of knock-out pistils with wild-type Col-0 pollen. Following a 2 hour pollination, the knock-out pistils were stained with aniline blue to visualize pollen grain adherence and pollen tubes. As shown in Figure 3, some knock-out lines show reduced pollen grain adherence and pollen tube penetration while others appear similar to wild-type Col-0 stigmas. Scale bar = 50μm

**Supplemental Fig. 2.** Col-0 pistils pollinated with *copi* mutant pollen grains. *α1-copi, β-copi allele 2*, and *γ-copi* pollen show either less pollen grain adherence or shorter pollen tubes when compared to the other *copi* mutant pollen grains. Scale bar = 50μm.

